# Translational buffering tunes gene expression in mice and humans

**DOI:** 10.1101/2025.05.16.654561

**Authors:** Shilpa Rao, Aden Y Le, Logan Persyn, Can Cenik

## Abstract

**Background:** Translational buffering refers to the regulation of ribosome occupancy to offset the effects of transcriptional variation. While previous studies have primarily investigated translational buffering in yeast under genetic variation or environmental stress, it remains unclear how widespread this is across mammalian genes in various cellular contexts.

**Results:** We performed a uniform analysis of 1,515 matched ribosome profiling and RNA-seq datasets from humans and mice. This resource enabled us to assess translational buffering through comparative analysis of variation in ribosome occupancy and RNA expression, and by examining the relationship between mRNA abundance and translation efficiency. We found that translational buffering is partly conserved between humans and mice; homologous genes showed moderate cross-species correlation in mRNA–translation efficiency relationships and strong enrichment of shared buffered genes, particularly those encoding ribosomal, RNA-binding, and proteasomal proteins. Although identified buffered genes associate with specific sequence features, these alone are insufficient to predict translational buffering, highlighting the importance of cellular context. Genes exhibiting translational buffering show lower variation in protein abundance in cancer cell lines and tissues. We also observed that translationally buffered genes are more likely to be haploinsufficient and triplosensitive, suggesting a demand for stringent dosage limits.

**Conclusions:** We hypothesize two models of translational buffering, namely the “differential accessibility model” and the “translation initiation rate model”, suggesting that different transcripts align with one or the other. Our study explores the translational buffering potential of genes across diverse conditions, elucidates their distinctive features, and provides insights into the mechanisms driving this effect.

## Background

mRNA variability arises from the inherent randomness of transcriptional events and in response to extrinsic fluctuations in the cellular environment. The randomness in transcription is driven by the stochastic nature of molecular interactions, whereas extrinsic noise reflects variation among cells in global factors such as metabolic state, signaling activity, cell cycle stage, and overall cell physiology [1–3]. In addition, genetic differences arising from aneuploidy, cell type diversity or those present within the natural population can further determine the amplitude of variation [4–7]. The effect of mRNA variation under these conditions is reflected at the protein levels for some genes. In other cases, however, it can be counteracted at the post-transcriptional level to minimize its effect on the protein abundance [2,4–9]. The mechanisms that can achieve this attenuation remain poorly characterized possibly involving excess protein degradation or translational buffering [4,6,10–15].

In this study, we define “translational buffering” as the post transcriptional mechanism wherein alterations in mRNA abundance is counteracted by changes in ribosome occupancy of transcripts. Early studies using allele-specific analyses of ribosome profiling and mRNA sequencing in yeast hybrids found that differences in mRNA abundance did not frequently lead to corresponding differences in ribosome occupancy [16,17]. More recent work profiled eight natural *S. cerevisiae* isolates using parallel RNA sequencing and ribosome profiling, and found that strain-to-strain transcriptional variation is buffered at the translational level [18]. While these studies focused on the impact of genetic differences on gene expression, translational buffering also underlies the cellular response to environmental stress. In yeast cells experiencing acute oxidative stress, aside from a small set of transcriptionally induced antioxidant genes, most mRNA expression changes were counteracted by opposite changes in ribosome occupancy resulting in negligible net protein change [19]. Similarly, translational buffering of certain genes have been reported in mammalian cells, specifically under estrogen receptor alpha depletion conditions [20]. Another comparative study that profiled both the transcriptome and translatome of three primary organs across five mammalian species and one bird observed buffering, characterized by reduced divergence at the translatome layer of especially essential and housekeeping genes [21]. Together, these studies of the impact of genetic variation, stress conditions and evolution suggest that cells often buffers or attenuates transcriptomic differences at the translation step, thereby stabilizing the proteome across different genotypes and environments.

Nonetheless, the prevalence of translational buffering among mammalian genes across different cellular contexts remains poorly characterized. Using matched measurements of mRNA abundance and ribosome occupancy from RNA-seq and ribosome profiling experiments, we can assess two related but distinct types of relationships between mRNA abundance and ribosome occupancy (see [10,15] for discussion of analogous comparisons in the context of mRNA-protein abundance relationship). First, in a given condition, we can determine the effect of mRNA abundance on ribosome occupancy across transcripts. This is informative to understand gene specific differences in the efficiencies of translation. Modest correlation between mRNA abundance and ribosome occupancy levels and a higher correlation between ribosome occupancy and their corresponding protein levels [22] iterates the significance of ribosome occupancy determinants in understanding the steady state levels of different proteins in a given condition. Second, ribosome profiling with matched mRNA-seq enables exploration of the effect of mRNA variation of a particular gene on its ribosome occupancy across different experimental conditions.

In this study, we analyzed the effect of mRNA variation on ribosome occupancy across diverse cellular conditions by leveraging a repository called RiboBase, harboring 1,282 human and 995 mouse ribosome profiling data with matched RNA-seq experiments [23]. The vastly increased number of experimental samples provides much higher statistical power to detect genes that are translationally buffered. Our definition of translational buffering is based on the observation that some genes show minimal variation in the ribosome occupancy levels despite having varied mRNA abundance across hundreds of experiments. This implies a negative correlation between the translation efficiency (TE) and mRNA abundance of buffered genes. Overall, our study identifies the potential of genes to be translationally buffered and elucidates the possible mechanisms and rationale for this phenomenon.

## Results

### Estimation of translation buffering potential of more than 8,000 genes

To systematically identify genes that exhibit translational buffering, we analyzed how variation in mRNA abundance affects ribosome occupancy and TE in over 8,000 genes (Methods). We leveraged 2,277 matched ribosome profiling and RNA-seq datasets from 90 human and 82 mouse tissues and cell lines compiled in a comprehensive database (RiboBase)[23]. Using this resource, we calculated TE similarly to [23], with some modifications. We estimate TE using a compositional regression model [23], rather than the commonly applied approach of using a logarithmic ratio of ribosome occupancy to mRNA abundance. The ratio-based approach introduces spurious correlations between the resulting TE and the mRNA abundance, which can confound interpretation [24,25]. In contrast to [23], we do not summarize TE by cell lines and instead use estimates from individual samples to capture a higher dynamic range of mRNA abundance which is essential for analysing translational buffering **(Additional file 2: Tables S1-S4)**. Next, we implemented a geometric Bayesian-multiplicative imputation procedure that preserves compositional structure and relative proportions, yielding more reliable TE estimates. Lastly, we introduced an additional quality control filter to exclude samples displaying non-linear relationships between mRNA abundance and ribosome occupancy (Methods). Such non-linearities often result from cellular perturbations, such as genetic modifications or pharmacological interventions and render regression residuals unreliable estimates of TE (Methods). In total, we retained TE estimates for 8433 human and 8135 mouse candidate transcripts from 965 human and 550 mouse samples (1515 total), respectively **(Additional file 2: Tables S5-S6)**.

For most genes, ribosome occupancy scales linearly or near-linearly with mRNA abundance, hence TE is only weakly dependent on transcript abundance (**Fig. 1A**). We reasoned that deviations from this pattern could be used to nominate genes exhibiting translation buffering. To identify such cases, we calculated pairwise correlations among TE, ribosome occupancy, and mRNA abundance across all human and mouse samples. The resulting wide distribution of correlation values suggests that mRNA variation differentially relates to TE and ribosome occupancy across genes **(Fig. 1B-C**, **Additional file 1: Fig. S1 A-B)**. We used strong negative correlation between mRNA abundance and TE as a primary indicator of translational buffering. As expected, there was considerable overlap between the set of genes with lowest correlation between either mRNA-TE or mRNA-Ribosome occupancy (176 out of top 250).

**Fig 1:**
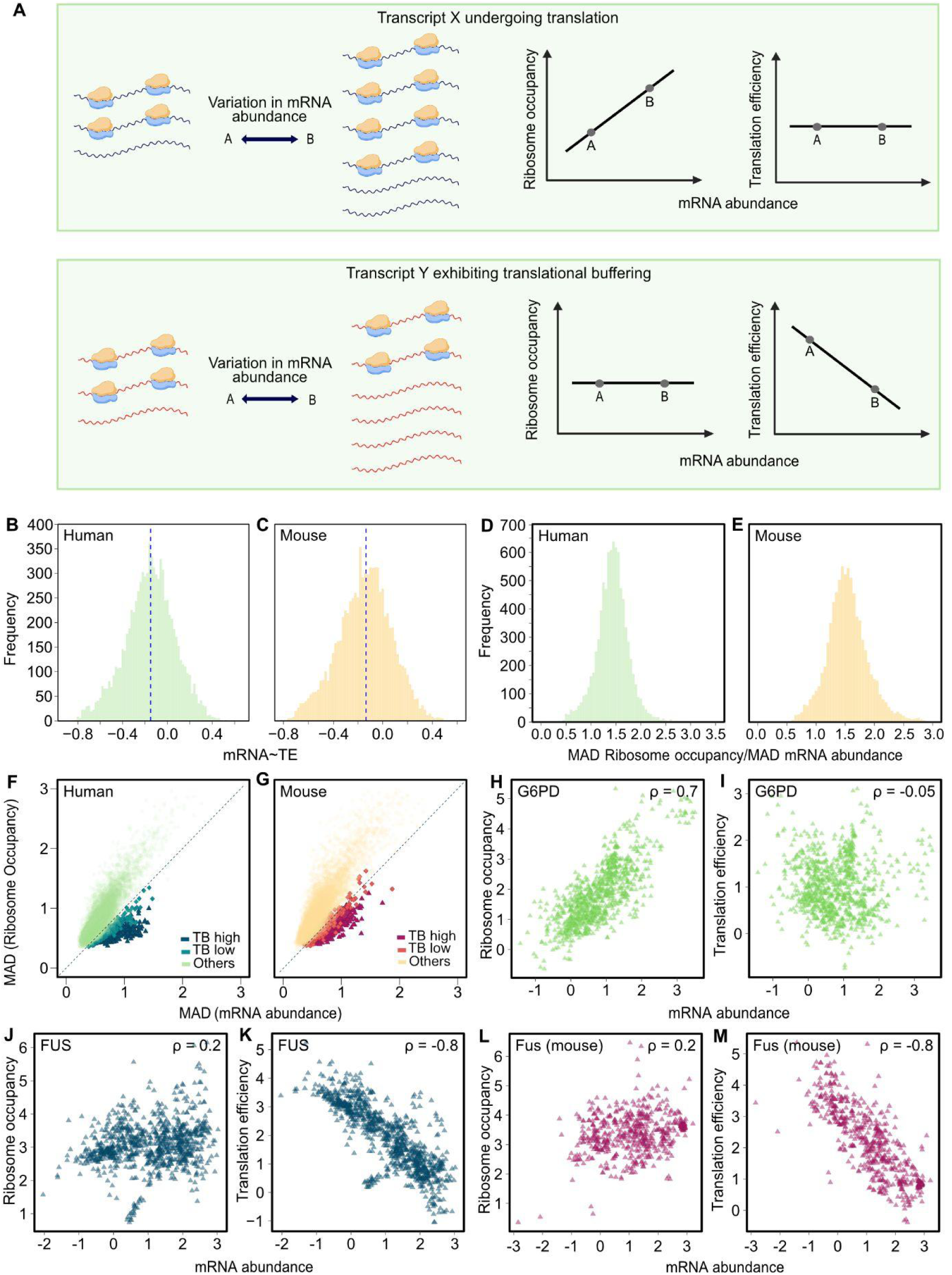
**A**. Schematic depiction of translational buffering: Transcript X shows linear relationship between mRNA abundance and ribosome occupancy resulting in no correlation between translation efficiency (TE) and mRNA abundance between condition A and B. Transcript Y shows no relationship between mRNA abundance and ribosome occupancy resulting in a negative correlation between translation efficiency and mRNA abundance between condition A and B. Hence transcript Y is considered to be translationally buffered. **B**. Distribution of Spearman correlation between mRNA abundance and translation efficiency calculated from ribosome profiling and matched RNA-seq of 965 human samples for 8433 human genes. The blue dashed line indicates the median of the correlation **C**. Distribution of Spearman correlation (⍴) between mRNA abundance and translation efficiency calculated from ribosome profiling and matched RNA-seq of 550 mouse samples of 8135 mouse genes. **D**. Distribution of the ratio of median absolute deviation (MAD) of ribosome occupancy vs MAD of mRNA abundance for human genes. **E**. for mouse genes. **F.** Scatter plot of MAD of mRNA abundance and MAD of ribosome occupancy for all genes in RiboBase for humans **G.** for mouse. Each point indicates a gene and the color depicts their buffering category. The dashed line indicates the line of equivalence**. H. & I.** Relationship between mRNA abundance and ribosome occupancy or translation efficiency of human non-buffered gene; G6PD, **J & K.** human buffered gene; FUS, **L & M.** mouse buffered gene; Fus. Each point is the information from a ribosome profiling and matched RNA-seq experiment for the given gene. All values are clr normalised (see methods)

As a second criterion, we identified genes with minimal changes in ribosome occupancy relative to variation in mRNA abundance. Our rationale is that if ribosome occupancy remains relatively constant despite substantial fluctuations in transcript abundance, this may indicate compensation at the level of translation. To quantify this, we calculated the median absolute deviation (MAD) for both ribosome occupancy and mRNA abundance across all samples in RiboBase **(Fig. 1D-E)**. We then determined genes with low ribosome occupancy MAD relative to their mRNA MAD **(Fig 1F-G)**. Based on these two criteria, genes were ranked and categorized as “TB high” (top 250), “TB low” (next 250), and “Others” (remaining genes) **(Additional file 2: Tables S7-S8).** Example genes in each category from human and mouse are shown in **Fig. 1H-M** and **Additional file 1: Fig. S1 C-H**.

### Conservation of translational buffering across species and its determinants

We hypothesized that if translational buffering is functionally important or inherently associated with specific transcript features, it would be conserved across species. To test this, we assessed the relationship between the mRNA-TE Spearman correlation values for homologous genes in human and mouse. We found a moderate correlation (∼0.6), indicating that the relationship between mRNA abundance and TE is partially conserved between these two species **(Fig. 2A)**.

**Fig. 2:**
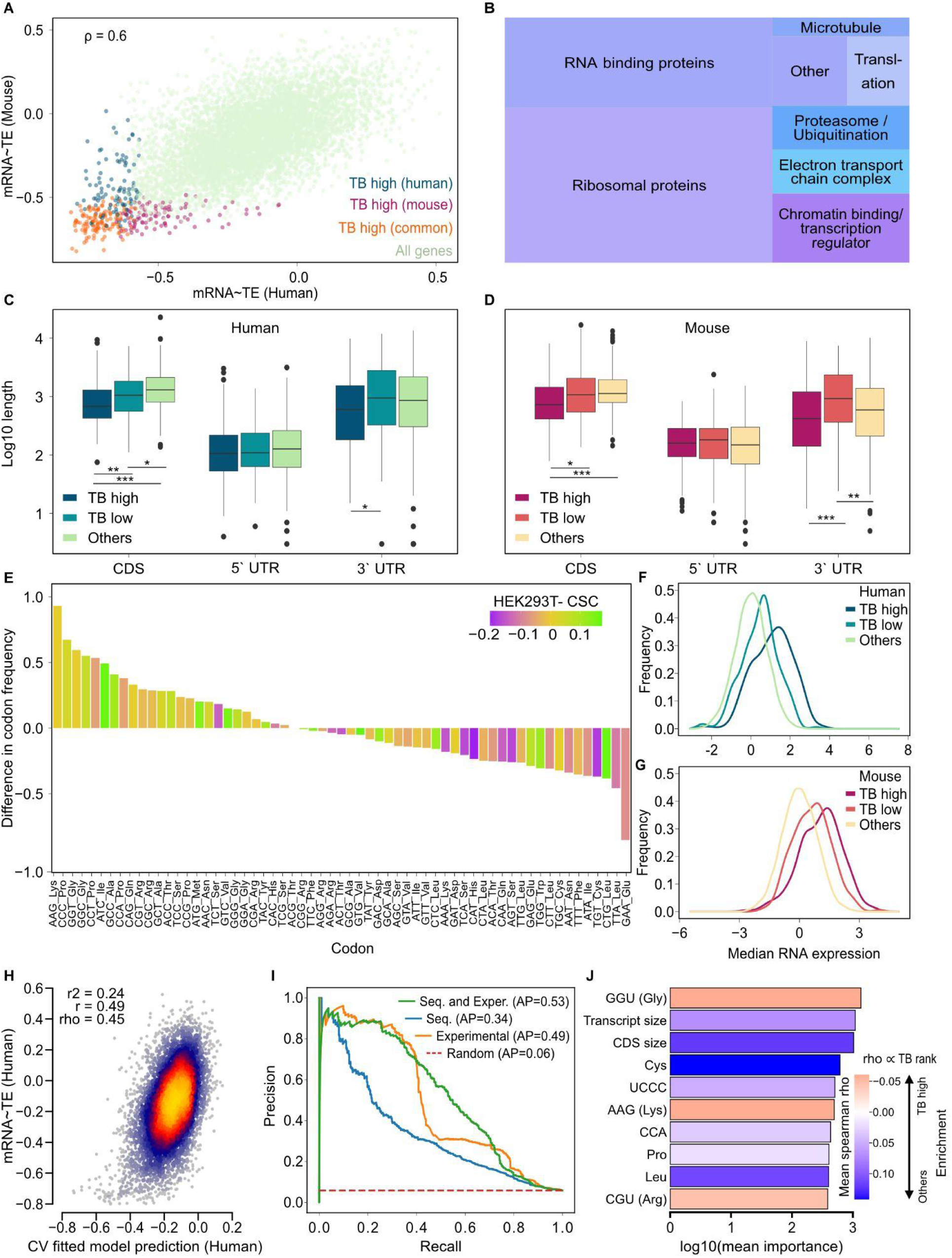
**A.** Scatter plot showing the relationship between mRNA abundance ∼ translation efficiency (TE) correlations between human and mouse (Spearman correlation coefficient = 0.6, p-value < 2.2e-16). **B.** Treemap showing the categories of genes that overlap between human and mouse top 250 (TB high) identified translationally buffered genes. Box plot showing the length of CDS, 5’-UTR and 3’-UTR of the three categories (TB high, low and others) of **C.** human genes (TB high n=250, TB low n= 245; Others n=495) **D.** mouse genes (TB high n=250, TB low n= 250; Others n=500) The y axis indicates log_10_ values. Gene in TB others category were selected after a matching procedure (See Methods). The * indicate p values. * < 1*10^-3^, ** < 1*10^-4^, *** < 1*10^-5^ **E.** Bar plot showing the difference in codon frequency between “high” and non-buffered genes The bars are colored according to the codon stabilization coefficient obtained from [29]. Density plot of median mRNA abundance across all samples in all three categories in **F.** human (TB high n=250, TB low n= 250; TB others n=7933) and **G.** mouse (TB high n=250, TB low n= 250; TB others n=7635). **H.** Scatter plot comparing predicted and observed human mRNA-TE Spearman correlation across 10 CV folds. The r^2^, Pearson (r), and Spearman (rho) correlation coefficients between predicted and observed are also shown. **I.** Precision-recall curves of LGBM models trained on three different sets of features: Sequence, Experimental, and both sets combined (Seq. and Exper.). Average precision (AP) for each model is shown along with the performance of randomly predicting class labels. **J.** The top 10 most important features determined by the sequence LGBM classification model (Methods). The bar color corresponds to the Spearman correlation between the feature and each gene’s translational buffering candidate rank. Note that a feature with a negative Spearman correlation indicates an enrichment in genes with lower mRNA–TE correlation. n refers to the number of genes.

Notably, 126 genes were shared among the top 250 translationally buffered genes (**Additional file 2: Table S9;** Fisher’s exact test odds ratio 83.4; p-value < 2.2*10^-16^). These genes include ribosomal proteins (n=55), RNA binding proteins (n=32), chromatin binding/transcriptional regulators (n=11), proteasomal complex proteins (n=7), genes involved in the electron transport chain complexes (n=6) and 15 others **(Fig. 2B)**. This distribution of genes suggests that genes encoding for components of macromolecular complexes may preferentially be enriched for translational buffering **(Additional file 1: Fig. S2A-B)**.

The conservation of translational buffering suggests that intrinsic transcript properties might contribute to this phenomenon. Hence, we compared various features, and found that the translationally buffered gene set had lower median CDS length in human and mouse (Wilcoxon rank-sum test p-value < 1.0 *10^-3^ comparing TB high and TB others; **Fig. 2C-D)**. Our analysis also reveals that highly ranked genes show associations with particular features, indicating an underlying hierarchy in translational buffering potential **(Additional file 1: Fig. S3A-B)**. Further, GC content of 5’-UTR was analysed for in both human and mouse with the later species showing a significantly lower 5’-UTR GC content potentially indicative of a propensity for increased secondary structures in the buffered set (Wilcoxon rank-sum test p-value 1.2*10^-7^ of TB high vs. TB others in mouse) **(Additional file 1: Fig. S3C-D, Additional file 2: Tables S10-S11)**.

Growing evidence suggests that translation initiation rates can be modulated by elongation speed, which is itself influenced by codon usage [26,27]. To understand if codon usage patterns are associated with translational buffering, we next analyzed codon properties across buffered and non-buffered human gene sets. Buffered gene candidates (both TB high and low) have a higher proportion of certain codons including that of lysine (AAG), proline (CCT, CCC, CCA), alanine (GCT), and glutamine **(Fig. 2E, Additional file 1: Fig. S4A, Additional file 2: Table S12)**.

The codon adaptation index quantifies how closely a gene’s codon usage aligns with that of highly expressed genes [28]. Genes in the buffered gene set had a higher codon adaptation index than the non-buffered set. Specifically, 28.4% of TB high genes, 14% of TB low genes and 9.3% of genes in the other category fall within the top decile (>90^th^ percentile) of codon adaptation index **(Additional file 1: Fig. S4B, Additional file 2: Table S13)**. Next, we analyzed the enrichment of certain codons associated with mRNA stability in HEK293T cells [29]. We observed a significant correlation between codon stabilization coefficient and the difference in codon frequency between buffered and non-buffered genes for both TB high (ρ =0.3, p-value= 0.02) and TB low (ρ =0.3, p-value= 0.01) categories, indicating that buffered genes are slightly enriched for codons associated with higher mRNA stability.

Given the observation with higher codon adaptation index and stability coefficients, we next analyzed if translationally buffered genes had higher average transcript abundance. We observed that the TB high transcripts had a slightly higher median mRNA abundance across all samples compared to genes with TB low and non-buffered gene sets (Wilcoxon rank-sum test p-value 6.9 * 10^-58^ for TB high vs. others and 1.2 * 10^-15^ for TB low vs. others; **Fig. 2F, Additional file 2: Table S7)**. Approximately, 52.8 % of TB high genes, 21.6% of TB low genes and 8.3% of genes in the other category fall within the top decile (>90th percentile) of median RNA expression. A similar trend was evident in mouse samples which predominantly represent primary tissue samples (Wilcoxon rank-sum test p-value 1.7*10^-51^ for TB high vs. others and 6.4*10^-23^ for TB low vs. others; **Fig. 2G, Additional file 2: Table S8)**. Although genes with the highest MAD values are not translationally buffered, we observed that non-buffered genes in both humans and mice exhibit lower variation in mRNA abundance across samples. These findings suggest that genes with either very high or very low MAD values are less likely to be translationally buffered **(Additional file 1: Fig. S4C-D)**. Collectively our analysis suggests that translationally buffered genes differ in intrinsic features, such as length, GC content, codon usage bias and expression levels.

To test if these sequence features are sufficient to determine translational buffering, we trained regression and classification models on features derived from the mRNA sequence and experimental data (Methods). These features were previously shown to be predictive of TE [30] and mRNA stability [31]. Specifically, we used light gradient-boosting machine (LGBM) models. The objective of the regression task was to predict the mRNA-TE Spearman correlation. An LGBM regression model trained only on sequence features achieved an r^2^ of 0.24 (**Fig 2H**). For the binary classification task, TB high and TB low genes were assigned the positive label and all other genes, the negative label. We trained models on three different sets of features: i) a model trained only on sequence features, ii) a model trained only on experimental features, and iii) a model trained on both combined feature sets (Methods). The models scored an average precision of 0.34, 0.49, and 0.53, respectively, while randomly assigning labels based on class balance achieves an average precision of 0.06 (**Fig. 2I**). The classification model trained on sequence features attributed high importance to many of the same features found to be enriched in TB genes discussed earlier (e.g. the codons GGU (Gly) and AAG (Lys), and CDS length; **Fig. 2J**). The enrichment of these codons or lower CDS length is associated with lower mRNA–TE correlation (negative spearman correlation) and hence a lower TB rank (rho ∝ TB rank). However, the modest performance observed suggests that the mRNA sequence alone is insufficient to predict translational buffering. The models that incorporate the experimental features outperform the sequence only model, further supporting the idea that cellular context is important in predicting translational buffering (**Fig. 2I**).

To investigate potential trans-acting determinants of translational buffering, we focused on 1,394 human RBPs as classified by [32]. Specifically, we examined correlations between the RNA expression of each RBP and the TE of all other genes across samples. This analysis revealed that the expression levels of many RBPs are significantly enriched for either positive or negative correlations with the TE of buffered genes (**Additional file 1: Fig. S5**). In particular, we note that RNA expression of many buffered RBPs is enriched for negative correlations with the TE of other buffered transcripts. These results suggest that, rather than considering translational buffering in isolation for each transcript, buffering effects may be coordinated at the translational level and influenced by shared trans-acting factors such as RBPs. Network-based approaches have been valuable for RNA co-expression and are only now being applied to TE covariation [33]. However, the correlative nature of these analyses limits causal inference. For example, although many ribosomal proteins appear to influence the buffering of other ribosomal proteins, they themselves may be regulated by a non-ribosomal RBP—so the apparent effects could reflect upstream regulatory influences. Together, the analysis suggests that both cis and trans acting factors might contribute to translational buffering of genes.

### Translational buffering offsets the impact of mRNA variation on protein abundance

Translational buffering could be expected to reduce fluctuations in their protein amounts due to changes in mRNA abundance **(Fig. 3A)**. To evaluate this, we compared the protein abundance variation using mass spectrometry data from 949 human cancer cell lines [34], 32 human primary tissues [35] and 41 mouse tissues [36]. Consistent with the expectation, buffered gene sets (TB high and low) had lower median absolute deviation in protein abundance compared to all non-buffered genes across all three datasets: in cancer cell lines (Wilcoxon rank-sum test p-value 5 *10^-39^ for TB high vs. others and 3 *10^-23^ for TB low vs. others) **(Fig 3B; Additional file 1: Fig. S6A)**, in human **(Fig. 3C; Additional file 1: Fig. S6B)** and mouse tissues **(Fig. 3D; Additional file 1: Fig. S6C).** To ensure that this reduced protein abundance variability was not simply explained by mRNA abundance differences, we selected non-buffered gene sets that are matched to buffered categories (both high and low) in terms of their mRNA MAD (Methods). Even after matching on variance in mRNA abundance, we observed a similarly lower deviation in protein abundance of buffered genes in human cancer cells (Wilcoxon rank-sum test p-value 2 *10^-29^ for TB high vs. others and 4.5 *10^-19^ for TB low vs. others) and mouse tissue samples (Wilcoxon rank-sum test p-value 8.4 *10^-26^ for TB high vs. others and 3 *10^-8^ for TB low vs. others). However, the difference in MAD (protein abundance) became statistically insignificant for human tissue samples **(Additional file 1: Fig. S6D-F)** as translationally buffered genes exhibited lower deviation in mRNA abundance compared to non-buffered genes in this particular dataset. These results indicate that translational buffering contributes to reduced protein variability across conditions, beyond what can be explained by mRNA variation alone.

**Fig. 3:**
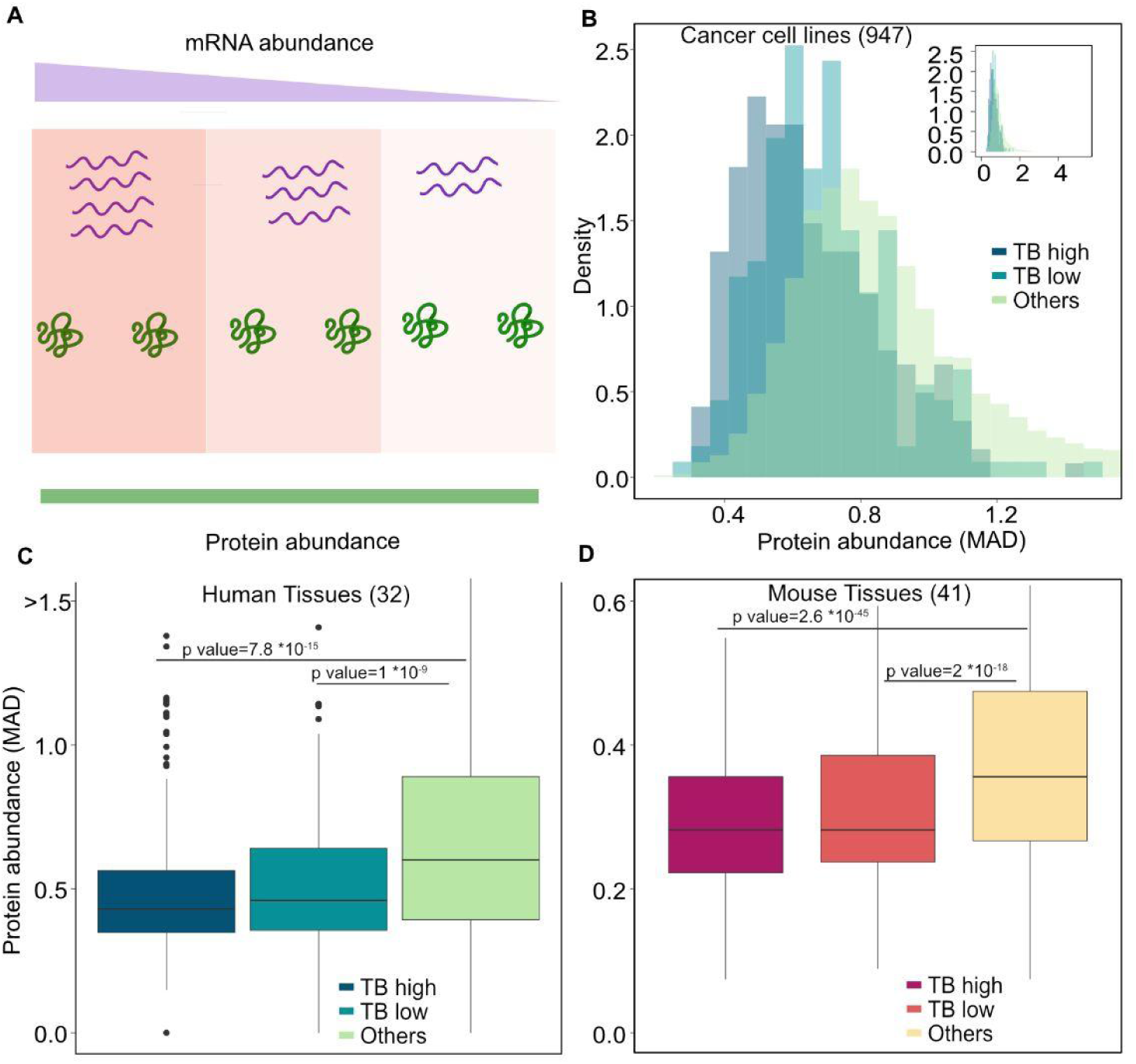
**A.** Schematic of the mRNA protein relationship in translational buffering **B.** Normalized histogram showing the MAD of protein abundance of genes in the three categories from human cancer cell lines (TB high n=221, TB low n= 202; Others n= 7954) **B.** Boxplot showing the MAD of protein abundance of genes in the three categories from **C.** human primary tissue (TB high n=233, TB low n= 227; Others n= 12167) and **D.** mouse tissues (TB high n=189 TB low n= 199; Others n= 6113). n refers to the number of genes

### Translational buffered genes are more likely to exhibit haploinsufficiency and triplosensitivity

Translational buffering dampens the impact of transcript abundance fluctuations on protein output and may serve as a mechanism to stabilize protein abundance across conditions. This property suggests the possibility that translational buffering preferentially occurs in dosage-sensitive genes as an evolved mechanism to maintain proteomic homeostasis. Alternatively, if buffering mitigates the consequences of reduction of transcript abundance, it may render genes more tolerant to dosage perturbations, predicting a depletion of haploinsufficient genes among the buffered set. To evaluate these competing hypotheses, we examined the correspondence between translational buffering and gene-level dosage sensitivity using population-based metrics **(Fig. 4A)**. Specifically, we first used the probability of loss of function intolerance (pLI score) as provided by the gNOMAD database. We found that buffered gene sets had higher pLI scores (median for TB high= 0.86, TB low= 0.76 and TB others= 0.0018), indicating increased likelihood of haploinsufficiency **(Fig. 4B)**. A chi-squared test revealed a highly significant difference in the distribution of genes across score ranges among the three categories (χ² = 519.5, df = 18, p < 2.2× 10⁻¹⁶), indicating that gene distributions differed substantially between them.

**Fig. 4:**
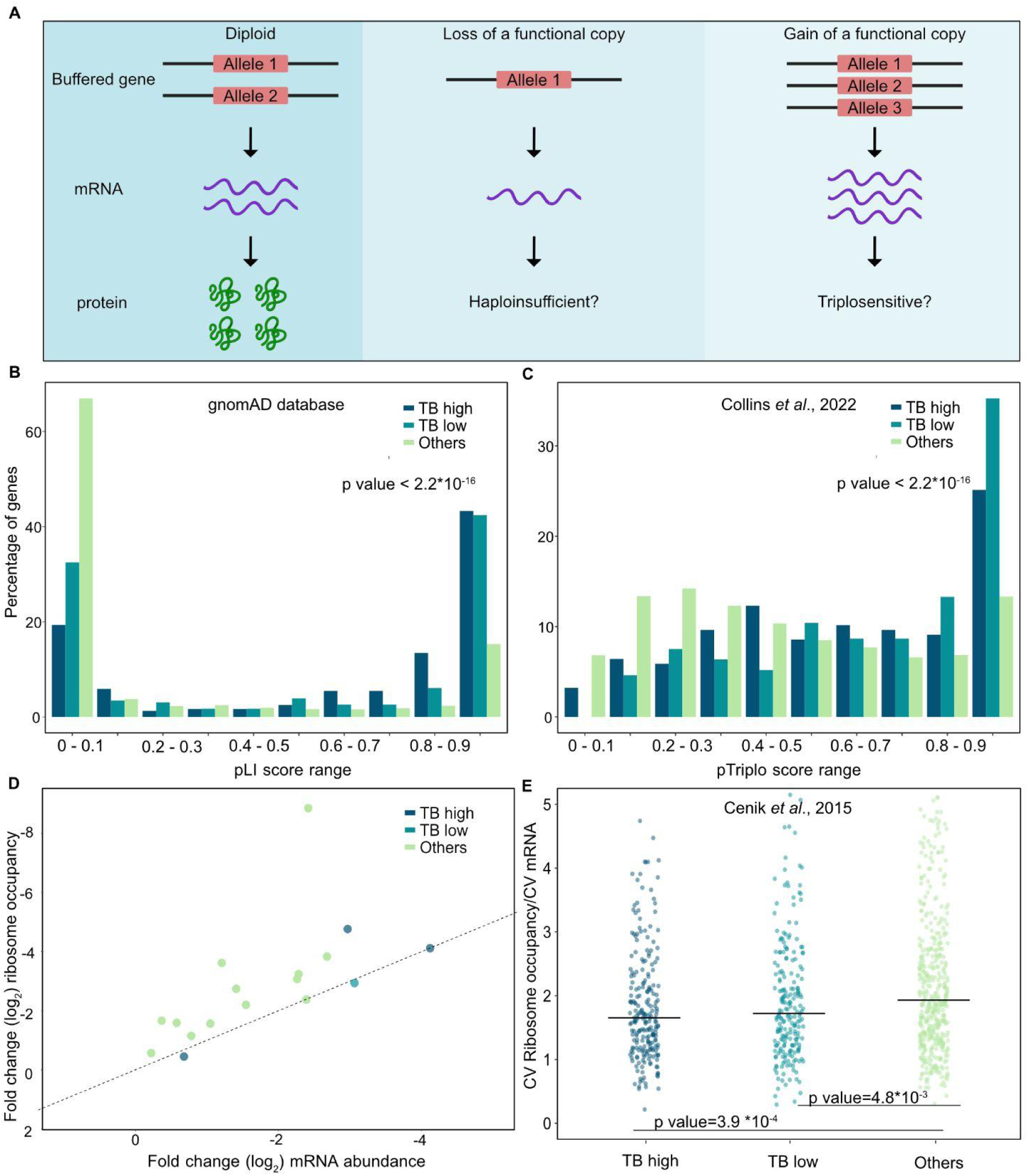
**A.** Schematic for the hypothesis whether the buffered gene sets are intolerant or tolerant towards dosage perturbations **B.** The gNOMAD database provides a pLI score (Probability of Loss of Function Intolerance) that predicts how likely a gene is able to tolerate loss-of-function mutations. Bar plot shows that buffered gene sets have a higher median pLI score (TB high n=238, TB mid n= 231; TB others; n= 18728) **C.** Collins et al. 2022 predicts the triplosensitivity for all autosomal genes referred to as pTriplo score. The bar plot shows a higher median pTriplo score (TB high n=187, TB mid n= 173; TB others; n= 11128) than the non-buffered set. n refers to the number of genes **D.** Plot showing the log2 fold change in mRNA abundance and ribosome occupancy in samples involving CRISPRi/siRNA knockdown of buffered and non-buffered genes from RiboBase. The color indicates the buffering category. Each dot represents a specific gene.Only genes that show significant change in mRNA abundance and Ribosome occupancy (p value <0.05) are plotted. **E.** Plot showing the ratio of Ribosome occupancy (CV) and mRNA abundance (CV) of genes analysed from lymphoblastoid cell lines. The genes in the “other” category were selected by a matching approach (Methods). The horizontal bar indicates the median of the ratio was lower in buffered genes than non-buffered genes (TB high n=235, TB mid n= 222; TB others; n= 457).

To further investigate the relationship between translational buffering and dosage sensitivity, we analyzed a second dataset that estimates haploinsufficiency and triplosensitivity for autosomal genes [37]. Genes in the translationally buffered sets had higher pHaplo scores (median for TB high= 0.802, TB mid= 0.697 and TB others= 0.48; χ²(18) = 114.8, p value= 4× 10⁻¹⁶, **Additional file 1: Fig. S7**) and pTriplo scores (median for TB high= 0.638, TB low= 0.785 and TB others= 0.43, χ²(18) = 151.3, p-value< 2.2× 10⁻¹⁶) compared to non-buffered genes (**Fig. 4C)**. These findings suggest that translationally buffered genes are more likely to exhibit dosage sensitivity and may have evolved for more stringent control of protein abundance.

The observed propensity for haploinsufficiency and triplosensitivity implies that translational buffering alone is insufficient to compensate for the large (∼50%) perturbation at the transcript level. Instead, translational buffering can likely counterbalance only minor fluctuations in mRNA abundance. This interpretation is also consistent with depletion of genes with high variability at the mRNA level among those that exhibit translational buffering **(Fig. 1F)**.

To further test this interpretation, we analyzed if experimental manipulation via siRNA or CRISPR knockdown could be compensated at the ribosome occupancy level (Methods). These perturbations, like haploinsufficiency and triplosensitivity, involve substantial reductions in transcript abundance and thus represent major perturbations beyond the range of typical physiological variation. For both TB high and TB low gene sets, experimental mRNA knockdown led to concordant reductions in ribosome occupancy proportional to the observed decrease in mRNA abundance **(Fig. 4D)**. However, the limited number of experiments with direct perturbation of buffered genes limit the generalizability of this observation.

Finally, we analyze the effect of natural variation at the level of RNA on ribosome occupancy levels [6] across lymphoblastoid cell lines from a diverse group of individuals. In contrast to the experimentally induced knockdowns or dosage sensitivity, this setting captures endogenous, inter-individual variability in transcript abundance under physiological conditions. As before, we selected genes with matched mRNA MAD across the three sets (TB high, low and others; **Fig. 4E, Additional file 2: Table S14**). Despite matched transcript-level variation, TB high genes exhibited significantly lower variability in ribosome occupancy compared to the non-buffered group (Wilcoxon rank-sum test p-value=3.9*10^-4^ and 4.8*10^-3^ for TB high or low vs. others, respectively). Taken together, these findings suggest that translational buffering likely functions as a homeostatic mechanism in response to physiological variation, but is insufficient to mitigate the effects of large-scale transcript perturbations.

### The association of translationally buffered genes with the translational machinery varies in response to changes in mRNA abundance

We next explored the potential mechanisms that may give rise to translational buffering. Specifically, we considered two non-mutually exclusive models by which mRNA abundance might be decoupled from ribosome occupancy. In the first, the “differential transcript accessibility model”, mRNA abundance determines the fraction of transcripts that are accessible to the translational pool. In this scenario, an increase in mRNA abundance would be accompanied by a proportionally smaller increase in the fraction of transcripts entering the translating pool for buffered genes, compared to non-buffered genes. In the second, the “initiation rate model”, the rate of translation initiation per transcript scales inversely with mRNA abundance. Under this model, as mRNA abundance increases, translation initiation on each transcript is reduced, thereby lowering the number of ribosomes per transcript. However, this mechanism allows a proportional increase in transcripts entering the translational pool for buffered genes, similar to non-buffered genes **(Fig. 5A)**.

**Fig. 5:**
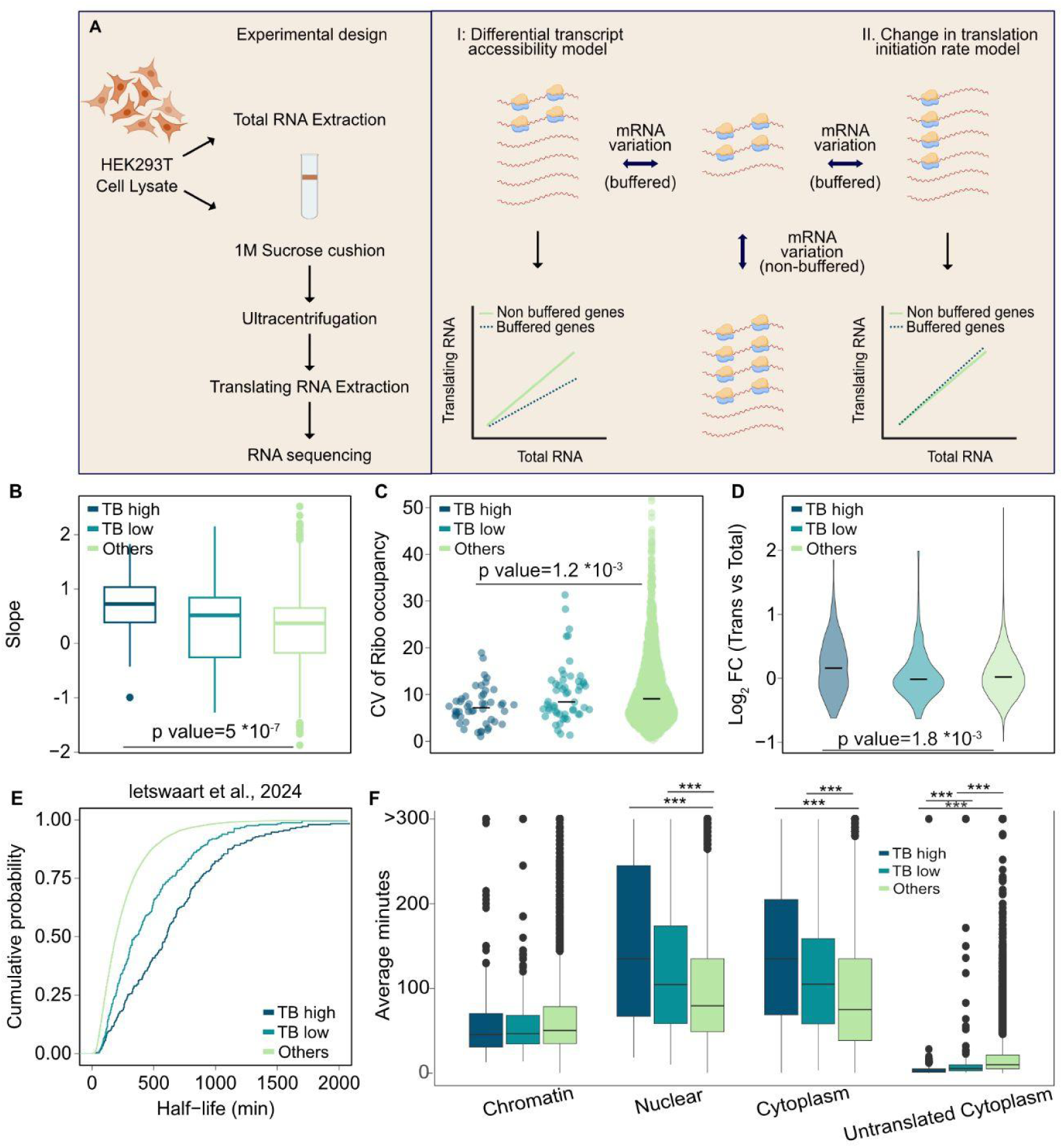
**A.** Schematic of experimental design to analyze the possible mechanism of translational buffering. **B.** Box plot showing the slope between translational RNA and total RNA (See Methods) for the buffered and non-buffered gene set (TB high n=48,TB low n= 52; TB others n=2157). The slope was higher in the buffered gene set which indicates that as the total mRNA abundance changes, a considerable fraction of buffered transcripts enter the translating pool as the non-buffered gene set. **C.** Sina plot showing the CV of ribosome occupancy levels of the three gene sets from the ribosome profiling experiment carried out in parallel to the RNA-seq of total and translating RNA (See Methods) (TB high n=48, TB low n= 52; TB others; n=2153). **D.** Violin plot showing the log_2_ fold change between the translational mRNA and total mRNA of all genes present in three categories (TB high n=250, TB low n= 250; TB others n=11889). **E.** Empirical cumulative distribution function (ECDF) plot showing the distribution of half life of mRNA for the three categories of gene sets (TB high n=249, TB low n= 246; TB others n=17670) **F.** Box plot showing the half life measurements of the three categories of gene sets (TB high n=249, TB low n= 246; TB others n=17670) in different compartments including the chromatin, nucleus, untranslated cytoplasm and the cytoplasm. The half life measurements of mRNA were obtained from [38]. n refers to the number of genes. For visualisation purposes, values greater than 300 have been capped at 300. The p value is after Bonferroni correction. * < 4*10^-3^, ** < 4*10^-4^, *** < 4*10^-5^

To evaluate the relative contributions of these models, we performed sequencing of both total and translating RNA fractions in HEK293T cells (Methods), leveraging variation in mRNA abundance across replicates. We considered genes that were in the top half in terms of mRNA variation as measured by coefficient of variation, and excluded genes with poor linearity between total and translating RNA (Methods). We then calculated the slope of the linear relationship between total and translating mRNA abundance across replicates. We observed a broad range of slopes (from < –1 to ∼1), where a slope of ∼0 would indicate minimal changes in the translating pool despite variation in total mRNA, and a slope near 1 would reflect a proportional increase in translation. Negative slopes, observed in some cases, likely reflect measurement errors and technical artifacts (e.g., ribosome dissociation during ultracentrifugation).

We found that genes in the “TB high” category exhibited significantly higher slopes compared to non-buffered genes (**Fig. 5B, Additional file 2: Table S15;** Wilcoxon rank-sum test p-value 5 *10^-7^ for TB high vs. others; 1.26 *10^-1^ for TB low vs. others). Yet, consistent with our earlier observation from RiboBase, these genes vary less at the ribosome occupancy levels (lower CV) compared to non-buffered gene set (**Fig. 5C**; Wilcoxon rank-sum test p-value 1.2 *10^-3^ for TB high vs. others; 7.5 *10^-1^ for TB low vs. others), indicating that increased access to the translational pool does not translate into proportional increases in ribosome loading. A similar trend was observed when genes from non-buffered set were selected to match the mRNA CV of the buffered gene set, confirming that buffered genes have a higher tendency to enter the translational pool despite comparable mRNA variability (Wilcoxon rank-sum test p-value 1.2 *10^-5^ for TB high vs. others; 1.3 *10^-1^ for TB low vs. others), while having lower ribosome occupancy variation (Wilcoxon rank-sum test p-value 1.8 *10^-2^ for TB high vs. others; 8.7 *10^-1^ for TB low vs. others **Additional file 1: Fig. S8A-B)**.

One caveat of this approach is the assumption that transcripts recovered following ultracentrifugation are exclusively associated with ribosomes. However, some transcripts may be bound to large ribonucleoprotein complexes that co-sediment with ribosome-associated mRNAs. Given this possibility, the slope for buffered transcripts could also be higher if their initiation rate does not change but the excess of buffered transcripts are bound by large ribonucleoprotein complexes.

To further examine the relationship between transcript abundance and engagement with the translational machinery, we employed an alternative analytical approach. We compared the relative abundance of transcripts in the translating pool versus the total RNA fraction between buffered and non-buffered gene sets. Consistent with the previous analyses, genes in the TB high category had a relatively higher fraction of genes in the translational pool compared to non-buffered genes as measured by calculating the log_2_ fold change between the translating pool and total RNA (Methods) (Wilcoxon rank-sum test p-value 7.8 *10^-8^ for TB high vs. others; **Fig. 5D, Additional file 2: Table S16)**. This enrichment could potentially be biased by differences in mRNA abundance. Even after matching for transcript abundance, buffered genes retained a significantly higher association with the translational pool (Wilcoxon rank-sum test p-value 1.3 *10^-29^ for TB high vs. others; 7.3 *10^-29^ for TB low vs. others; **Additional file 1: Fig. S8C**) indicating that mRNA abundance does not alone account for their preferential engagement with the translation machinery. Taken together, these findings suggest that both the initiation rate models and the differential transcript accessibility may contribute to translational buffering. Further studies will be required to clarify the relative contributions of these mechanisms and delineate their molecular mechanisms.

Finally, we analysed mRNA stability of the three categories. We compared the half life of buffered vs non-buffered transcripts [38]. Translationally buffered genes had longer mRNA half-life compared to non-buffered genes (median: TB high; 625 minutes, TB low; 352.5 and TB others; 195 minutes) (Wilcoxon rank-sum test p-value 2 *10^-60^ for TB high vs. others; 2.1 *10^-25^ for TB low vs. others; **Fig. 5E**). This observation is consistent with genes in the TB high category being enriched for codons associated with higher stability (**Fig. 2E**). To gain insight into the kinetics of half life of transcripts in each of the subcellular locations, we leveraged TimeLap-seq data [38]. This analysis suggested that the transcripts from the TB high category spend more time in the cytoplasm (half life in cytoplasm; median: TB high; 135 min, TB low; 105 min, others; 75 min, Wilcoxon rank-sum test p-value 3.4 *10^-18^ for TB high vs. others; 4.9 *10^-9^ for TB low vs. others) however are quicker in loading to the polysome fraction (half life in untranslated cytoplasm; median: TB high; 2.3 min, TB low; 5.4 min others; 10 min; **Fig. 5F**, Wilcoxon rank-sum test p-value 2.3 *10^-72^ for TB high vs. others; 9.4 *10^-21^ for TB low vs. others). Translationally buffered transcripts also seemed to be exported more slowly from the nucleus than non-buffered transcripts (half life in nucleoplasm; median: TB high; 135 min, TB low; 106 min, others; 80 min, Wilcoxon rank-sum test p-value 3.5 *10^-8^ for TB high vs. others; 1.8 *10^-5^ for TB low vs. others). On comparing the subcellular half life of transcripts similar in their whole cell half life (Methods), the buffered transcripts (TB high) have a higher half life in nucleus (half life in nucleus; median: TB high; 135 min, others; 115 min) **(Additional file 1: Fig. S8D)**. Overall, these observations indicate that buffered transcripts differ from non-buffered sets in their RNA flow dynamics and have a higher possibility of association probably for a longer time with the translational machinery than the non-buffered transcripts.

## Discussion

Ribosome occupancy of certain transcripts remains relatively unchanged, even when their mRNA levels vary across conditions. This phenomenon has been referred to as “post-transcriptional buffering” in several yeast studies [17–19] and as “translational buffering” in others [20,39]. Across these studies, differences in species evaluated, statistical methods and definitions of buffering have resulted in conflicting interpretations [10,15,25,40].

To systematically characterize translational buffering at the transcriptome scale and to evaluate this phenomenon across different conditions, we leveraged a compendium (RiboBase) of more than a thousand ribosome profiling datasets with matched RNA-seq measurements [23]. This resource enabled us to quantify how variation in mRNA abundance across samples influences ribosome occupancy and TE. The breadth of experimental conditions, encompassing a range of human cell lines and mouse primary tissues, provided robust statistical power to detect translational buffering across diverse cellular contexts. Though we relied on a subjective threshold for defining genes as “TB high or low”, we emphasize that genes span a continuum of translational buffering potential, with certain genes demonstrating more stringent control (TB high and low) over ribosome occupancy levels than others (TB others). This was further supported by testing different thresholds to define TB-high and TB-low groups. Higher cutoffs made the three categories more similar across the characteristics assessed as shown in **Additional file 1: Fig. S9**

Estimating the translational efficiency from ribosome profiling and matched RNA-seq data remains an area of debate. The method we used to calculate TE is based on a compositional regression model [23], rather than the commonly applied approach of using a logarithmic ratio of ribosome occupancy to mRNA abundance. The ratio-based approach introduces spurious correlations between the resulting TE and the mRNA abundance, which can confound interpretation [24,25]. Several methods that have been developed previously for measuring changes in TE [41–43]. These are designed to be used in experimental designs where there is a treatment or manipulation that is assayed along with controls. While these methods are highly valuable in assessing differential TE, they are unable to accommodate the type of meta-analyses described in our study. The combination of a large sample size across diverse conditions independently evaluated in two species, along with our TE calculation method, offers a distinct advantage for studying translational buffering and expanding its interpretation across species.

Our analyses revealed substantial overlap in translationally buffered genes between human and mouse, despite the differing sources of the underlying datasets. While human samples were predominantly derived from cell lines, mouse samples originated largely from tissues. This concordance across species suggests that translational buffering may be conserved for a subset of genes. Many of these genes encode components of multiprotein complexes, such as ribosomes, the electron transport chain, and the proteasome, all of which have previously been shown to exhibit translational buffering in yeast as well [17–19]. However, the set of translationally buffered genes can vary depending on cellular or environmental context, as with observations in natural isolates and different growth conditions [18].

In both human and mouse datasets, translationally buffered genes are typically highly expressed and often encode dosage-sensitive complexes, including the ribosome. The change in absolute transcript abundance for these highly expressed genes in response to intrinsic or extrinsic noise would be substantial compared to lowly expressed genes. This likely necessitates more stringent regulation. Hence, translational buffering may serve as a compensatory mechanism, limiting protein production from these abundant transcripts to ensure that protein synthesis remains within physiological limits.

In addition to their expression levels, transcripts also exhibit varying degrees of fluctuation across samples. Certain buffered genes show a higher stringency in their mRNA variation. For example, we observed that constitutively expressed genes like ribosomal genes and GAPDH exhibit modest variation in mRNA levels across conditions, consistent with findings in yeast [18]. Recent studies have also reported sub-Poissonian mRNA distributions for some constitutively expressed genes in yeast [44]. In contrast, other buffered gene candidates, such as FUS, display significant variation in mRNA levels across samples, suggesting less stringent transcriptional control. This may require a more robust buffering mechanism. However, whether these two cases differ in their buffering mechanisms remains to be determined. Additionally, it would be counterproductive to allow high variation in transcript levels only to later balance it at the translational level. While this could be due to stochastic fluctuations in mRNA, it is interesting to speculate that mRNA variability in certain genes may sequester regulatory molecules such as RNA binding proteins.

Many transcripts in the TB high category encode ribosomal proteins characterized by the presence of TOP motifs whose TE is known to be co-regulated in response to environmental cues mainly through the mTOR pathway [45]. Our analysis suggests that their TE is possibly altered in relation to fluctuations in transcript levels to potentially minimize variation in protein abundance. Recent reports suggest that TOP transcripts are known to be associated with sub-polysomal fractions and represent translationally repressed transcripts [46,47]. Such a repression may facilitate translational buffering in some transcripts.

Although we observed associations between specific sequence features and translational buffering, these are unlikely to be the sole determinants. This is supported by our machine learning model, which shows that sequence alone is insufficient to accurately predict genes exhibiting translational buffering. Furthermore, the model indicates that features within the coding sequence are more informative than those in the untranslated regions. The limited predictive power of sequence features alone suggests that cellular context likely plays a significant role in determining the translational buffering capacity. This is in contrast to translation efficiency itself which can be much more accurately predicted solely from mRNA sequence [30]

Beyond delineating the determinants of translational buffering, our study elucidates the adaptive value of this phenomenon. Earlier studies with a Drosophila brain tumor model suggested that fold change in protein abundance of genes involved in energy metabolism, transcription and translation have lower correlation with fold changes at mRNA levels (ρ= 0.23, ρ= 0.16, ρ= 0.45 respectively) than other complexes [48]. Other studies using aneuploidy cell lines indicated that though the mRNA expression correlates to the increased chromosome copy numbers, approximately a quarter of the proteins are less abundant than expected and more similar to their disomic context [8]. Our analysis suggests that regulation of ribosome occupancy by translational buffering could be one of the strategies adopted by the cells to achieve lower variation at protein level. Another rationale for some genes to exhibit translational buffering could be that cells could tune expression of genes regulated by the same transcription factors, thus allowing some targets but not others to ultimately end up with higher or lower protein levels.

The mechanistic basis of translational buffering remains incompletely understood. Recent work has shown that upstream open reading frames can buffer main ORF by inducing ribosome stalling, particularly through terminal diproline motifs under conditions of limited ribosome availability [49]. Our RNA-seq experiments of translating versus total RNA fractions provides evidence supporting the feasibility of two models: differential transcript accessibility to the translational machinery and modulation of translation initiation rate particularly in response to changes in mRNA abundance. The accessibility model is supported by analyses of transcript half-lives in nuclear and cytoplasmic compartments, which suggest that transcript availability to ribosomes may be regulated at the level of subcellular localization. In contrast, the fast loading of buffered transcripts on the polysomes also argues for the initiation rate modulation model. Since the half-life measurements reflect steady-state levels, investigating how mRNA variation influences the spatial and temporal dynamics of transcripts could provide valuable insights into the mechanisms underlying translational buffering.

While this study expands our understanding of translational buffering, it has several important limitations. Firstly, our study is limited to genes that are robustly expressed in most samples thereby missing information on lowly or cell-type specifically expressed genes. Genes that are possibly translationally buffered in a particular cellular-context or tissue specific manner would escape our identification criteria. Further, our analysis relies on steady-state measurements of RNA abundance and ribosome occupancy, which may fail to capture dynamic or transient regulatory mechanisms that contribute to translational buffering. Next, most of the reported analyses are correlative in nature, therefore experimental manipulation of mRNA variability to analyse the effect on ribosome occupancy/TE might be needed. A potential drawback in understanding the models of translational buffering in response to natural variation in mRNA abundance is distinguishing true signal from technical noise. This challenge, coupled with a low replicate number (four), complicates the interpretation of slopes and the prediction of hypothesized models. Incomplete discrimination between mechanistic models, could potentially be due to insufficient molecular details. For instance, it remains unclear what specific molecular mechanisms are capable of sensing mRNA abundance variability and subsequently modulating translation initiation or accessibility. Clarifying this connection is crucial for distinguishing between models of translational buffering. We also note that ribosome occupancy is not always a reliable proxy for protein output, as additional factors such as translation elongation rates, ribosome stalling, and co-translational degradation can significantly influence the efficiency and outcome of protein synthesis.

Despite these caveats our study fills a gap in knowledge regarding translational buffering of genes in humans and mouse. Other model organisms such as *C.elegans* (nematodes), Drosophila, zebrafish have orders of magnitude less data making it difficult to assess translational buffering with high confidence. However, as more matched transcriptomic and translatomic datasets become available in other model organisms, extending this analysis will provide a broader perspective particularly in determining whether translational buffering is intrinsically encoded or primarily driven by cellular context. The availability of these datasets will also make it feasible to analyze translational buffering at the interspecies level. For example, many of the mitochondrial proteins were suggested to be translationally buffered under specific genetic backgrounds in *C.elegans* [50]. Hence, we speculate that this phenomenon may be conserved beyond mammalian genomes. Finally, complementary approaches such as live-cell imaging of translation at the single-transcript level under variable mRNA conditions could yield mechanistic insights beyond what bulk analyses allow.

## Conclusions

Overall, our work expands the catalog of buffered genes, reveals their evolutionary conservation, highlights their potential functional constraints (e.g., dosage sensitivity), and offers a conceptual and computational framework for studying translational buffering in broader biological settings.

## Methods

### Calculation of translational efficiency

The method for calculating translation efficiency (TE) is adapted from [23], with some modifications. In this study, TE is calculated for each gene for individual samples in RiboBase rather than summarized at the cell line level as in [23]. For this, matched RNA-seq (PCR-deduplicated) and ribosome profiling read counts (non-deduplicated and winsorised) for 19,736 genes in 1,078 human and 21,569 genes in 846 mouse experiments were used **(Additional file 2: Table S1-S4)**. The method of obtaining the RPF and RNA-seq read counts are described in [23]. We determined genes which have <1 count per million in more than 20% of the samples. In addition, we identified transcripts that lack a poly-A tail have artifactually “low” RNA -seq counts in experiments that rely on oligo-dT selection for library preparation, though their ribosome occupancy reads are well captured [6]. The corresponding raw read counts of all these transcripts were then collapsed and assigned to a “dummy gene”. This was necessary because these genes, though part of the composition in the sample, were excluded from downstream analyses given their low expression across the compendium. In total, the dummy gene captures read counts corresponding to 10,859 genes. A small number of samples wherein the “dummy gene” contributed to more than 70% of the total counts were removed from analysis, reducing the number of experiments to ∼1000.

A particular challenge in compositional data analysis is the handling of missing values. In this study, we used a geometric bayesian multiplicative imputation method from the R package zCompositions[51]. This function by default removes any samples which have more than 80% missing values in the sample (therefore removed 2 more samples). As the imputation method was parallelized, the resulting imputed values may differ slightly compared to a non-parallel implementation. Genes with imputed values were used for conversion to their centred log ratio (clr) values. To calculate the TE, clr-normalized counts were converted to isometric log ratios. These values were used to generate a line of best fit between the two parameters and the residuals were defined as TE for each gene in a given sample. Samples which had R^2^ less than 0.2 were removed as the residuals calculated for these samples could be unreliable. After these calculations, mRNA abundance, translational efficiency and ribosome occupancy values of 8,433 human genes from 965 samples and 8,135 mouse genes from 550 samples were retained for further analyses.

### Assignment of translational buffering scores

For each gene, Spearman correlation between mRNA abundance and either ribosome occupancy or TE was calculated across samples separately for human or mouse. Similarly, the median absolute deviation (MAD) was computed for mRNA abundance and ribosome occupancy for each gene and the ratio of MAD for ribosome occupancy to that of mRNA abundance was calculated. We then ranked genes based on two metrics: (i) mRNA-TE correlation (with the most negative correlation ranked highest) (ii) MAD ratio (with lower ratio ranked higher). We then summed the ranks from both metrics and genes were re-ranked, with lower sums corresponding to higher final ranks.The top 250 genes were categorised as “TB high”, the next 250 “TB low” and the remaining genes as “Others”. To assess across species conservation, Spearman correlation was also computed between the mRNA-TE correlations of human and mouse orthologs and genes were annotated according to their respective translational buffering categories.

### Gene ontology enrichment analysis

Lists of TB high genes in human or mouse were analyzed using the enrichGO() function, with the organism-specific annotation database (org.Hs.eg.db for human or org.Mm.eg.db for mouse) as reference using clusterProfiler R package [52]. Gene identifiers were supplied as gene symbols, and all genes in the current study were used as the background universe. Enrichment was carried out for Biological Process, with significance assessed by the hypergeometric test. P-values were adjusted for multiple testing using the Benjamini–Hochberg method, and terms with an adjusted p-value < 0.05 were considered significant. To reduce redundancy among highly overlapping GO terms, results were post-processed with the simplify() function in clusterProfiler, which merges semantically similar terms. Visualization of the non-redundant enriched GO terms was performed using dot plots (dotplot()) showing the top terms ranked by adjusted p-value.

### Sequence feature analysis

Transcript region lengths were extracted from the APPRIS annotations using custom shell scripts. Corresponding nucleotide sequences were retrieved with bedtools v2.29.1 [53] and nucleotide content was computed using the Bioconductor Biostrings package v2.54.0 [54]. For both humans and mouse, the log10-transformed lengths of coding sequence, 5’UTR and 3’UTR length and GC content for each region were calculated for all genes.

To control for confounding sequence features, non-buffered genes (Others) were matched to the buffered genes based on either CDS length (for comparing UTR lengths) or UTR lengths (for comparing CDS lengths) using the MatchIT package (nearest neighbor method) [55]. Similarly, for GC content comparisons, non-buffered genes were matched either on UTR GC content (for CDS GC content comparison) or on CDS GC content (for UTR GC content comparisons). Statistical differences between gene groups across sequence features were assessed using the Wilcoxon rank-sum test.

To analyze codon bias, codon frequencies were calculated for each human gene using custom R scripts and the BioString package [54]. Median mRNA abundance was computed across samples for each gene and the top fifth percentile of genes by expression was used to define the highly expressed set. The codon adaptation index was calculated using the cubar package [56]. Codon frequencies were computed for each buffering category and the difference in frequency between buffered (TB high & low) and non-buffered set (TB others) was calculated for each codon. To assess if the codon usage differences relate to transcript stability, the codon stabilization coefficient (CSE) obtained from [29] was compared to codon frequency difference using Pearson correlation.

### Sequence based predictive modeling of translational buffering

The sequence features used in the machine learning tasks were: i) the length, nucleotide composition, and {2-6}-mer frequencies across the 5’-UTR, CDS, 3’-UTR, and full transcript; ii) codon and amino acid usage; iii) wobble base composition; iv) the nucleotides at the -3, -2, -1, +4, and +5 Kozak positions; v) frequencies of dicodons that affect TE in yeast [57]; and vi) multiple secondary structure measures. The sequence features were computed with the same methods used previously [30]. The experimental features include various measures such as the number CLIP-seq and RIP-seq peaks for different RBPs [31]. LGBM models (lightgbm v3.2.1) [58] were trained with 10-fold cross validation. In the classification task, 20% of the training data for each fold was held as a validation set to calibrate the classifier. The default hyperparameters for the LGBM model were used except with the importance type set to “gain”. Feature importance was measured as the sum total information gain across all LGBM tree splits using that feature, averaged across all training folds.

### RNA binding protein (RBP) correlation analysis

To explore potential trans-acting determinants of translational buffering, we focused on 1,394 human RBPs as classified by [32]. We examined correlations between the RNA expression of each RBP and the TE of other genes across samples. P-values were corrected using the Bonferroni procedure, and Pearson correlations greater than 0.5 or less than -0.5 that met the p-value cutoff were considered significant for each RBP-gene pair. Subsequently, for each RBP, we performed Fisher’s exact test to evaluate whether significant correlations were enriched among buffered versus non-buffered genes. The Bonferroni procedure was also applied to determine the threshold for the odds ratio.

### Proteomics data and analyses

Protein abundance from 949 human cell lines [34], 32 human primary tissues [35] and 41 mouse tissues [36] were used to evaluate the effect of translational buffering on protein abundance variability. For each dataset, MAD of protein abundance was calculated across samples for each gene. To control for differences in transcript expression, non-buffered genes (TB others) were matched to buffered genes based on MAD of mRNA abundance using the nearest neighbor method implemented in the MatchIT package [55]. Differences in protein abundance variability between buffered and non-buffered gene sets were assessed using the Wilcoxon rank-sum test.

### Dosage sensitivity among translationally buffered genes

The gNOMAD database was used to retrieve the pLI score of all genes [59] followed by calculating the percentage of genes in each pLI score range. A second dataset providing predictions of haploinsufficiency (pHaplo score) and triplosensitivity (pTriplo score) for all autosomal genes [37] was used to assess the distribution of these scores across buffered and non-buffered gene sets. Comparisons were made to evaluate whether translationally buffered genes are enriched for dosage-sensitive characteristics. Differences between distributions were assessed using Pearson’s chi-squared test of independence.

### Differential expression analysis of knockdown experiments

Ribosome profiling experiments and matched RNA-seq involving siRNA/CRISPRi-mediated knockdown of genes categorized as *TB-high*, *TB-low*, or *others* were retrieved from the RiboBase database[23]. For each experiment, the RNA-seq and ribosome profiling counts were normalized using the TMM normalization [60]. Transcript specific dispersion estimates were calculated and differentially expressed genes in each study were identified using edgeR [61].

### Lymphoblastoid cell line data analysis

To assess the impact of natural variation in mRNA levels on ribosome occupancy, we analyzed lymphoblastoid cell lines from genetically diverse individuals using data from Cenik et al [6]. Matched RNA-seq and ribosome profiling count files were obtained from GEO (accession: GSE65912). Only samples with both RNA and ribosome occupancy data were included in downstream analyses. For each gene, the coefficient of variation (CV) was calculated separately for mRNA abundance and ribosome occupancy across individuals. The ratio of CV for ribosome occupancy to CV for mRNA abundance was then computed in R. To generate a background gene set (*TB others*), genes were matched to the *TB-high* and *TB-low* groups based on their mRNA CVs using the nearest-neighbor method in the MatchIt package [55].

### Cell culture

HEK293T cell lines were obtained from ATCC. HEK293T cell lines were maintained in Dulbecco’s modified Eagle’s media (DMEM, GIBCO) supplemented with 10% Fetal Bovine Serum (FBS, GeminiBio) and 1% Penicillin and Streptomycin (GIBCO, Life Technologies) at 37°C in 5% CO_2_ atmosphere. Cell lines were tested for mycoplasma contamination every 6 months and were consistently found negative.

### Translating and total RNA isolation and sequencing

Four replicates of ∼8 X10^6^ HEK293T cells were plated in 150 cm2 plates (see Supplementary text). After ∼18 h, cells were washed and then scraped with ice cold PBS, followed by centrifugation at 300g for 5 min. 600 μl Lysis buffer (20 mM Tris, 150 mM NaCl, 5 mM MgCl2, 1mM DTT, Triton and 100 μg/ml of cycloheximide) was added to the cell pellet followed by incubation on ice for 10 min and the lysate was clarified by centrifugation at 1300g for 10 min at 4°C. The cleared lysate (50 μl) was directly used for total RNA extraction by adding 700 μl of Trizol (Zymo Research). 200 μl of the lysate (translating RNA) along with 20 μl of the Ribonucleoside Vanadyl Complex was layered on the 1M sucrose cushion and subjected to ultracentrifugation at 38,000 rpm for 2.5 h at 4°C to isolate the translating RNA. The remaining 350 μl was used for ribosome profiling. The pellet was dislodged in 700 μl of Qiazol and stored -80°C until use the next day. Total RNA and translating RNA was extracted using Zymo RNA miniprep kit as per manufacturer’s protocol (Zymo Research). The concentration was estimated using nanodrop and the integrity of the extracted RNA was determined by 2100 Bioanalyzer (RNA 6000 Pico Kit, Agilient). 1μg of the extracted RNA was used for poly(A) selection using NEBNext poly(A) mRNA magnetic isolation kit (NEB, E7490S). This was followed by library preparation using NEBNext Ultra II RNA library prep kit for Illumina (NEB, E7770S) as per the manufacturer’s protocol using 7 PCR cycles. The quality and concentration of the libraries formed were determined by Bioanalyzer (High sensitivity DNA assay kit, Agilient). Equimolar amounts of the libraries were pooled and sequencing on NovaSeqX Plus (2X150 paired end).

### Ribosome profiling

For ribosome profiling, 350 μl of lysate was digested with 5 μl of RNase (Ambion) for 30 min at room temperature, layered on 1M sucrose solution followed by ultracentrifugation. The RNA was isolated using miRNeasy Mini kit and size-selected by running 1 µg of each sample in a 15% polyacrylamide TBE-UREA gel. 26–34 nt RNA fragments were excised, RNA fragments were extracted and precipitated as described in [62]. Ribosome profiling libraries were prepared using the D-Plex Small RNA-Seq Kit (Diagenode) as previously described [62]. The cDNA was amplified for eleven cycles. Each library was cleaned with AMPure XP beads and eluted with 30 μL of RNase-free water. To completely remove the primer-dimers, size-selection was done using a 12% polyacrylamide gel followed by excision of the fragment (∼200bp). The DNA was extracted by overnight incubation in a buffer (10 mM magnesium acetate tetrahydrate, 0.5 M ammonium acetate, 1 mM EDTA (pH 8.0) at room temperature followed by ethanol precipitation (-20°C, overnight) to recover the size selected libraries. The resulting libraries were analyzed with the Agilent High Sensitivity DNA Kit (Agilent). Each library was mixed in equimolar proportion for sequencing with the NovaSeqX Plus (2X150 paired end).

### Analysis of translating RNA/total RNA and ribosome profiling data

Ribosome profiling and the matched RNA-seq experiments (4 replicates each) were processed using RiboFlow [63] as before [62] and leveraged RiboGraph for data visualization [64]. Genes with >10 counts per million in more than 6 samples were retained. We then fitted a linear regression line using lm function in R and determined the slope between the translating and total RNA. Genes with an R² value less than the median, indicating a non-linear relationship between translating RNA and total RNA, were removed from. In addition, genes with low variability between replicates (i.e CV of RNA between replicates, < median CV) were also filtered out. After applying these, 2,257 genes were analysed. To ensure that differences in slope and the variation in ribosome occupancy were not due to differences in mRNA variation, genes in the non-buffered gene set were matched to the buffered gene set based on mRNA CV using the nearest-neighbor method in the MatchIt package [55]. To analyze the differential engagement of buffered and non-buffered gene sets with the translating pool, differential analysis between the translating and total RNA was performed using edgeR [61]. To ensure that resulting log2 fold change between the translating pool and total RNA, is not contributed by differences in mRNA abundance, genes in the non-buffered set were matched to the buffered gene set based on mRNA abundance (cpm) using the nearest-neighbor method in the MatchIt package [55].

### Subcellular mRNA half life analysis

mRNA half-life data were obtained from [38]. For each gene, we calculated the average mRNA half-life by taking the mean of replicate values across the chromatin, nuclear, cytoplasmic, and untranslated cytoplasmic fractions, along with the whole-cell mRNA half-life. This average was used as a representative measure of mRNA half life across subcellular compartments.

## Availability of Data and Materials

Both the raw data and processed sequencing file (ribo file) are available in the Gene Expression Omnibus (GEO) repository, GSE297442 [65] The source code used in the study is available at https://github.com/CenikLab/Translational-buffering/tree/Translational-Buffering under the GNU General Public License v3 [66]. In addition, the R source code used in the study is also available on Zenodo at DOI 10.5281/zenodo.18393475 [67].

## Acknowledgements

We thank Dr. Vighnesh Ghatpande for valuable suggestions. We used two LLMs; ChatGPT and UT Spark for the following purposes: 1. to improve clarity and grammar of the text and 2. To generate code for data visualization and statistical analysis which was subsequently reviewed, edited and tested before being implemented for final analysis.

## Peer review information

This article was peer reviewed at Review Commons and the original paper [68], reviewer reports [69,70] and point-by-point response [71] are available online.

## Authors’ Contributions

S.R, and C.C. wrote the manuscript. L.P developed the LGBM model, S.R. and A.L. carried out all other analyses; S.R. carried out all experiments. C.C. provided study oversight, conceptualized the study and acquired funding. All authors read and approved the final manuscript.

## Ethics declarations

## Ethics approval and consent to participate

Not applicable

## Consent for publication

Not applicable

## Additional files

Additional file 1: Fig S1-S9. Supplementary figures S1 to S9

Additional file 2: Tables S1-S16 Supplementary tables S1 to S16

